# Modeling disease progression in newly diagnosed type 2 diabetes

**DOI:** 10.1101/2020.05.04.076133

**Authors:** Kun Hao, Yanguang Cao

## Abstract

Type 2 diabetes (T2DM) is a progressive disease, which is primarily characterized by a decline in β-cell function and worsening of insulin resistance. Unfortunately, most interventions (lifestyle, diet, and therapeutic agents) for T2DM only provide a transient restoration of β-cell function and the progression is inevitable once it starts. To understand the natural progression of T2DM, a mechanistic model was developed to quantitatively characterize the dynamic interactions among β-cell function, plasma fasting glucose (PFG), fasting insulin (FI), and the degree of insulin resistance, starting from an early stage of T2DM over up to 8 years. The model was validated using clinical data to optimize the disease parameters. The restoration and deterioration rates of β-cell function were both predicted as 84.5 %/year and 1.10 /year for early stages of T2DM. The model predicted a positive correlation between the initial level of β-cell function at diagnosis and its maximum restoration potential, underscoring the importance of early diagnosis and intervention. After the treatment, β-cell function could be temporarily restored within several months, which has a long-term benefit in glycemic control. The maximal tolerated PFG level that permits β-cell function restoration was predicted to be around 8.33 nM; and the temporal restoration of β-cell function would be unlikely at a PFG level above this threshold. The intrinsic deterioration rates of β-cell function and insulin resistance were both critical factors for long-term glycemic control. In conclusion, our model provides a quantitative analysis of the natural disease progression in T2DM and yields insights into factors that are critical for long-term glycemic control.

## 1. Introduction

The β-cell dysfunction (BF) and insulin resistance are two primary pathophysiological factors for type 2 diabetes mellitus (T2DM) (Weyer et al., 1999). In the pre-T2DM stage, hyperglycemia-induced by insulin resistance is not apparent due to the compensational increase in insulin secretion. The compensational secretion by β-cells leads to the gradual loss of β-cell function. When β-cells fail to produce sufficient insulin to compensate for insulin resistance, plasma glucose would go up and BF would inevitably decline until overt T2DM. The natural history of T2DM has been demonstrated in several previous studies (Weyer et al., 1999; Kahn, 2007), most of which concluded that the onset and progression of T2DM were dictated mainly by the dynamics of β-cell deterioration, despite the different interventions and the varying patient population in these studies (Bagust and Beale, 2003). Once diagnosed, the initial control of hyperglycemia only temporarily (about one year) preserves BF to a minimal degree; most conventional treatments eventually fail to maintain BF and halt diabetes progression (Kahn, 2001). A natural deterioration process of BF, independent of glycemic control and insulin sensitivity, is believed to account for the many treatment failures in T2DM (Matveyenko and Butler, 2006; Tan et al., 2005).

Characterizing the disease progression with mathematical models is being frequently applied to quantitatively understand interactive pathological factors (Bies et al., 2008). T2DM is a polygenic and progressive metabolic disorder. Quantitatively assessing the progression of T2DM is beneficial to elucidating its complicated pathology and designing effective treatments. Several mathematical models have been developed based on the major pathophysiological defects in T2DM. De Gaetano et al. developed a mechanism-based model that involved several intrinsic risk factors in accounting for both the insulin sensitivity index (ISI) and BF and this model was primarily for simulation purposes (De et al., 2008). Topp et al. developed a model considering the mutual interactions of insulin, glucose, and BF (Topp et al., 2000). Ribbing et al. implemented this model in clinical settings for a short period of characterization of disease progression (about 1.5 years) (Ribbing et al., 2010). Two logistic functions were employed to describe the gradual change of insulin resistance and β-cell dysfunction, which were additionally applied to incorporate clinical treatment effects (Ribbing et al., 2010).

In the present study, we developed a diabetes disease model by integrating four major diabetic factors (glucose, insulin, BF, ISI) over a long period (up to 8 years after diagnosis). Data from several clinical trials were applied to validate the model in different conditions. The model provided quantitative assessments of the intrinsic β-cell deterioration and temporary restoration, the magnitude of glucotoxicity, as well as their long-term effect in the control of T2DM. Our quantitative analysis provided insights into diabetic disease progression, which are valuable for developing effective strategies in the long-term management of T2DM.

## 2. Materials and methods

### 2.1. Data sources and analytical method

The clinical data that reported the temporal profiles of PFG, FI, and BF were extracted from the literature (Tan et al., 2005; Alvarsson et al., 2008; Kahn SE et al., 2006; Levy J et al., 1998; UKPDS 33; UKPDS 16; UKPDS 13). The average curves were digitized for our analysis. All the studies were conducted in the newly diagnosed T2DM evaluating different interventions, including lifestyle change (conventional diet modification plus exercise), metformin, sulfonylureas, insulin, or thiazolidinedione. When the ISI and BF were not reported, we utilized the Homeostasis Model Assessment-1 (HOMA-1) to calculate the ISI and BF based on the values of PFG and FI at each time point (Levy JC et al., 1998). In total, 29 sets of profiles were obtained, **Fig. 1** depicts these profiles. As expected, there was a temporary restoration of BF after the intervention, which could last for about 1-2 years. Shortly after the temporary restoration, BF progressively declined, resulting in a gradual increase of FPG.

**Fig. 1.**
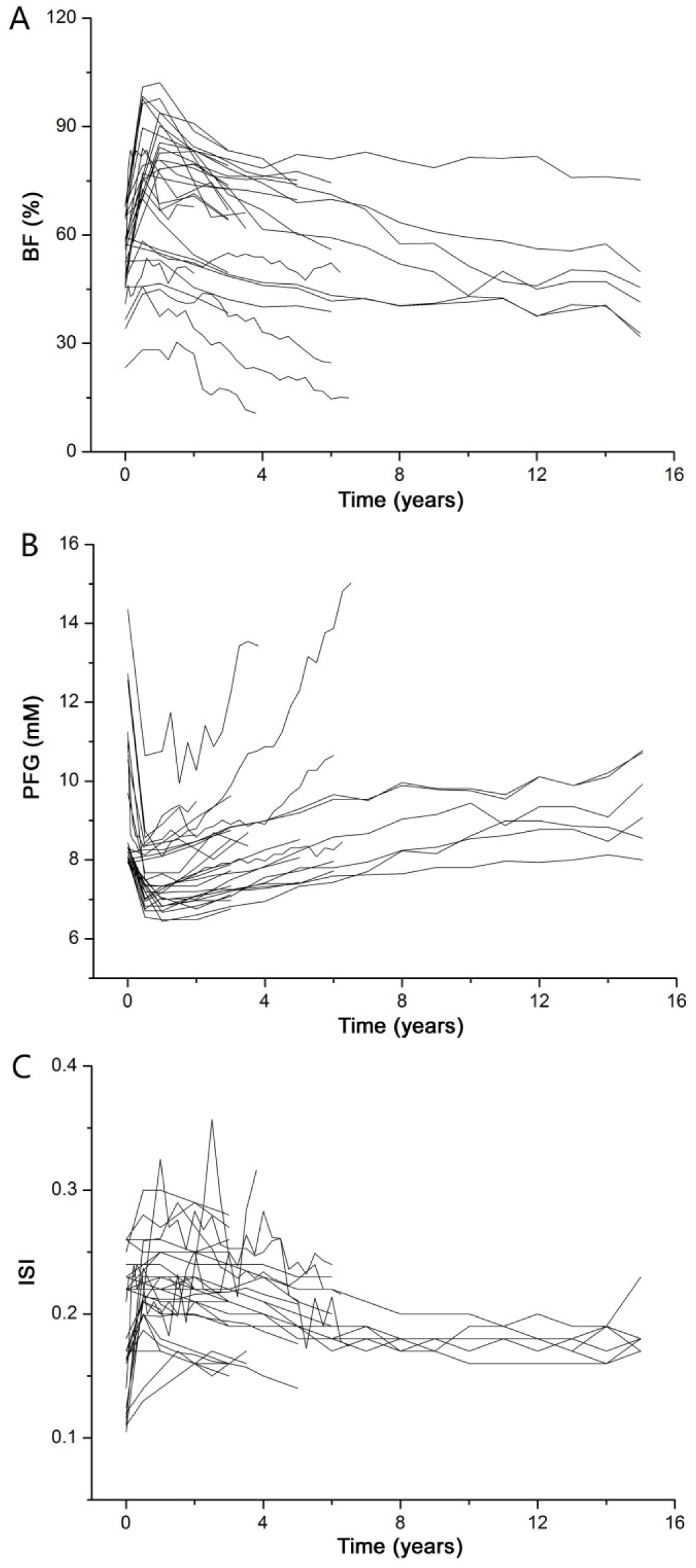
Time profiles of BF, PFG, and ISI from all clinical trials. BF and ISI are derived using the HOMA-1 model.

### 2.2. Diabetes Progression Model

The diabetic disease progression model is shown in **Fig. 2**. In the model, BF and PFG are considered as two dependent variables in diabetic progression, which are described by a physiological turnover model with zero-order input and first-order dissipation (Dayneka NL et al., 1993). The increase of PFG is a function of FI, drug treatment effect, and ISF. The differential equations are:

**Fig. 2.**
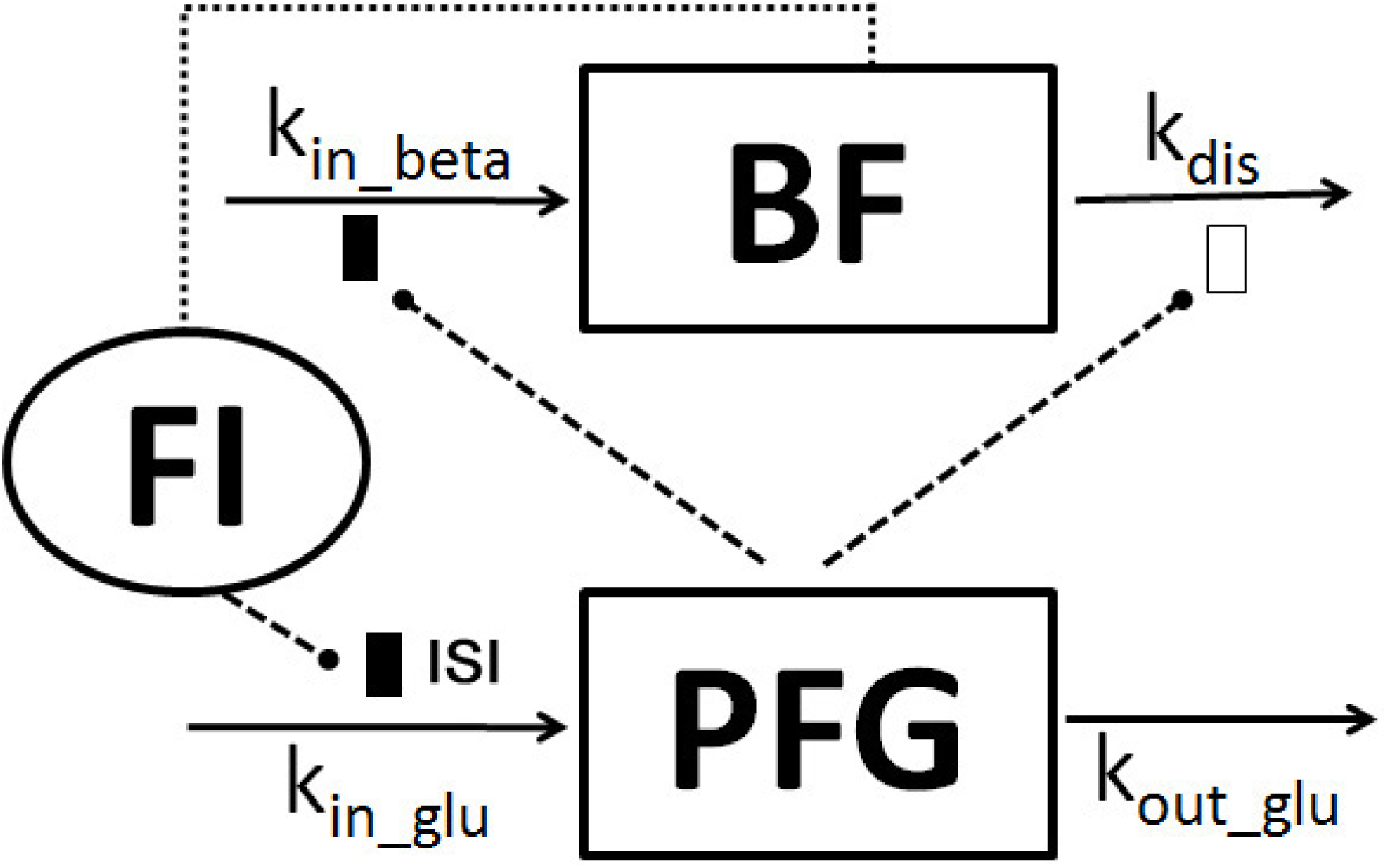
The structure of the diabetic disease progression model. Symbols are defined in the text. Lines with arrows indicate the turnover of the stated factors. Dashed lines ending in closed circles indicate the connected factors exert an action. The black box indicates an inhibition effect on β-cell restoration and plasma glucose production; the white box indicates an enhanced impact on β-cell deterioration. Dotted lines indicate the interaction between BF and FI.

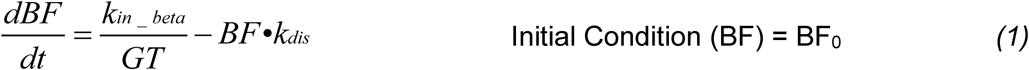

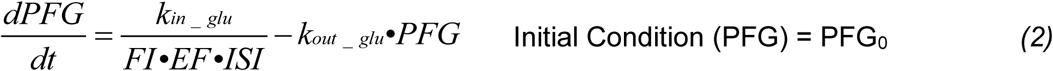

where BF is derived from the HOMA-1 model with normal BF defined as 100 %. BF is assumed to be controlled by two processes: the restoration of functional insulin-producing β-cells (*k*_*in_beta*_) and the decrease in functional β-cells (*k*_*dis*_) (Prentki M and Nolan CJ, 2006). Of note, *k*_*in_beta*_ and *k*_*dis*_ denote the apparent rate constants for BF gain and deterioration, which include changes in β-cell mass and functionality. The parameters *GT* and *k*_*dis*_ indicate glucotoxicity (equation 3) and β-cell intrinsic deterioration rate (equation 4). The parameters *k*_*in_glu*_ and *k*_*out_glu*_ are defined as the gluconeogenesis rate and the glucose utilization rate. *EF* represents drug treatment effect. Of note, in the HOMA-1 model, the treatment effects can potentially influence the calculations of BF and ISI. But the diverse mechanisms of interventions prohibit a specific definition of treatment effect. Therefore, a generalized term, *EF*, was defined to account for the effect of various treatments — for instance, the acute insulin-tropic effect by sulfonylureas and the inhibitive effect on gluconeogenesis by metformin. For insulin sensitizers such as thiazolidinedione and a restricted diet, the effects are reflected in ISI.

In the fasting state, insulin primarily acts as an inhibitor on endogenous glucose production in the liver with a limited effect on glucose utilization. This assumption was typically made at the fasting state in previous models (Gao W et al., 2011). Glucotoxicity is one of the major mechanisms that cause β-cell dysfunction and demise and is expressed using a function of PFG:

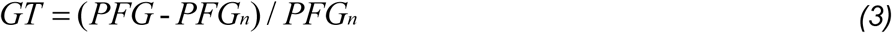

where PFG_n_ refers to normal PFG (4.5 mM), PFG that below and equal 4.5 mM shows no glucotoxicity. Besides glucotoxicity effect, BF also follows an intrinsic deterioration process which is described by:

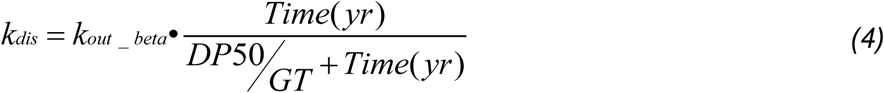

where *k*_*out_beta*_ is the deterioration rate of BF. *DP*_*50*_ is the time required to reach 50 % of the maximum deterioration rate. *Time (yr)* is the duration of time since diagnosis. It should be noted that *GT* is incorporated through both an inhibitive effect on β-cell restoration and an enhanced effect on β-cell deterioration, shortening the time required to reach 50 % of the maximum deterioration rate.

The physiological turnover of insulin is not defined as a dependent variable considering the fact that insulin turnover is very fast (in min) in comparison with the much slower process of diabetic progression (in years). The data would not support an accurate estimation of insulin turnover rate. Therefore, FI was derived from the HOMA-1 model based on reported PFG and BF at each time point, which was treated as an independent variable:

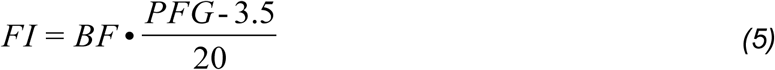

In the HOMA-1 model, an underpinning relation is FI = 5 × (PFG - 3.5) when BF = 100. The steady-state concentrations are: 5 mU/L for FI and 4.5 mM for PFG in healthy subjects (Matthews DR et al., 1985). When substituting these values into equation 2 with EF = ISI = 1 for healthy subjects, we obtained:

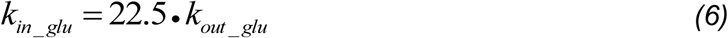

### 2.3. Statistical Analysis

The model was applied to analyze the literature-extracted data using a nonlinear generalized mixed effect modeling approach (ADAPT 5, Biomedical Simulations Resource, University of Southern California, Los Angeles, CA) (D’Argenio DZ et al., 2009). The model’s predictive performance was evaluated by a visual predictive check with 1,000 Monte Carlo simulations. Competing models were compared by their objective function, predictive performance, parameter variability, and residual errors. Glucotoxicity was simulated at several hyperglycemic conditions. The time courses of BF deterioration in different PFG conditions were summarized and compared to quantitatively assess glucotoxicity. Additional simulations were performed to evaluate the influence of β-cell deterioration rate and ISI on long-term glycemic control. The probabilities of PFG over 10 mM, 12 mM, and 14 mM and the probabilities of BF below 10 %, 30 %, and 50 % were chosen to reflect diabetic progression and disease severity.

## 3. Results

**Fig.1** summarized all datasets reporting the temporal profiles of BF, PFG, and ISI that were applied for model validation. As noted, the β-cells had already declined to 50 ∼ 60 % of the normal function at diagnosis in most patients. After intervention, BF increased briefly to a limited extent and then inevitably declined. The elevated PFG decreased temporarily after initiating diabetic treatments, and shortly returned to a rising trend (**Fig. 1B**). The nadir in BF profiles occurred concurrently with the peak in PFG profiles, which is in line with previous observations that β-cell dysfunction is the primary driving factor that dictates diabetic progression (Kahn SE 2001). At around 1.5 years after diagnosis, most interventions failed to further restore BF and patients started losing control of climbing PFG.

Insulin resistance is another pathological factor that contributes to diabetic progression, which had almost reached the most severe level at diagnosis. Insulin resistance typically occurs 5 ∼ 10 years earlier than the manifestation of diabetic symptoms (Weyer C et al., 1999). Insulin resistance did not progress much after diagnosis (**Fig.1C**). ISI profiles stay around 0.2 throughout the process of study for most patients.

As shown in **Fig. 3**, the developed disease model parameter of all 29 datasets sufficiently captured the temporal profiles of BF and PFG over a period of 8 years after diagnosis of T2DM. No systemic bias was noticed in model fittings. In addition to the trend, the model also well captured the inter-study variability, and the predicted percentiles largely overlapped with observations. The predicted PFG showed a lower limit around 6 mM, which is 0.5 mM higher than our assumption on the normal PFG in Equation 6, suggesting most patients rarely reached a normal level of PFG, regardless of interventions. Overall, both the trend and variability in the progressive trajectories of PFG and BF were both well predicted using the developed model.

**Fig. 3.**
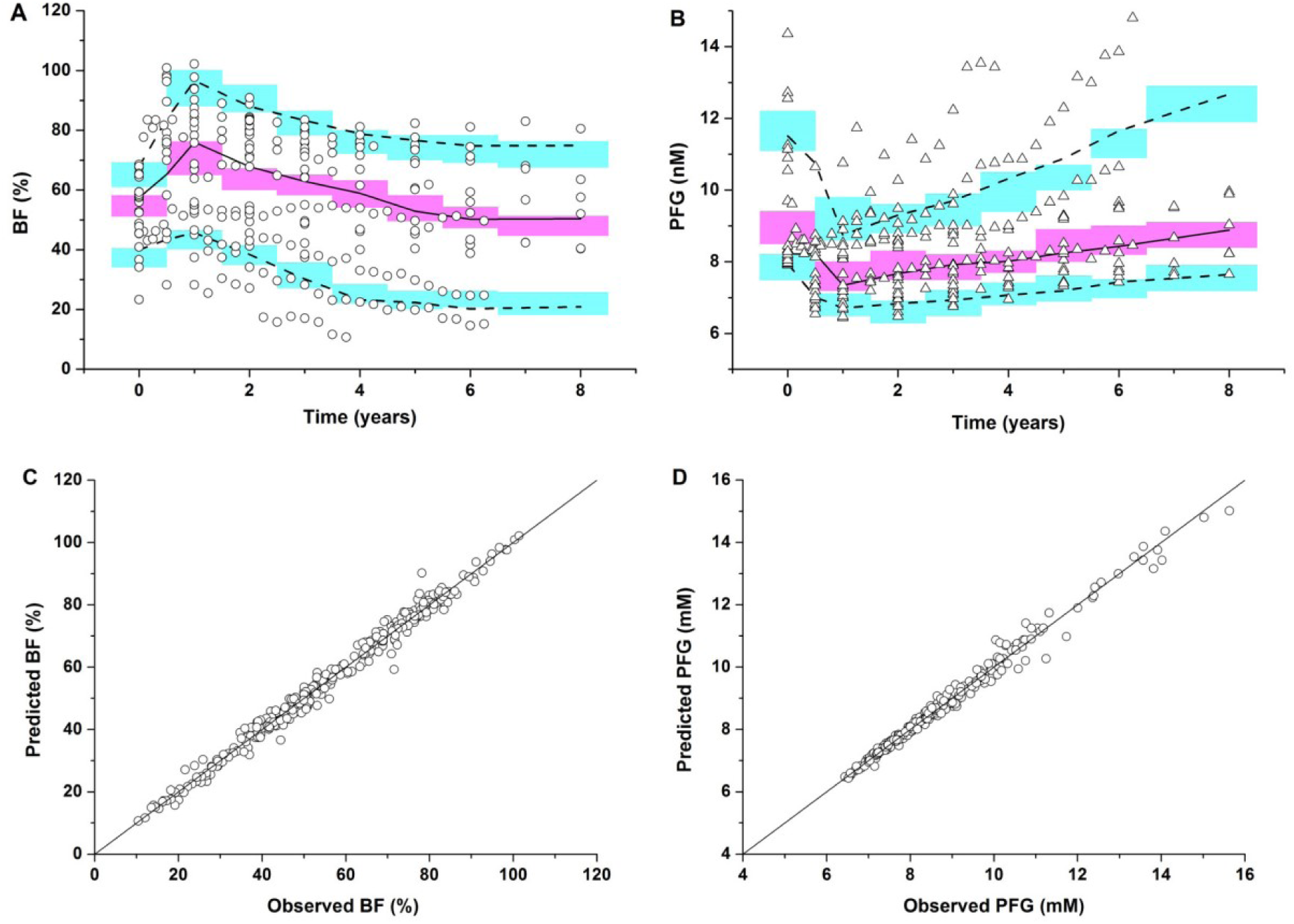
Visual Predictive check based on 1,000 simulations for BF and PFG over eight years. Legend: circle = real BF observations; triangle = real PFG observations; solid and dashed lines = the 50th, 10th and 90th percentiles for all real observations in each bin; red and green shaded areas = the 95% CI the 50th, 10th and 90th percentiles of the simulated data (**Fig. 3A and 3B**). The goodness of population fitting for BF and PFG (**Fig. 3C and 3D**).

The estimated parameters are summarized in **Table 1**. The deterioration rate for BF was predicted to be 1.10/year, which is close to the maximum deterioration rate at uncontrolled hyperglycemia. The restoration rate of BF is about 84.5 %/year, which indicates that β-cells have the potential to fully recover within a year when glucose is strictly controlled at the normal level and the intrinsic deterioration of BF is halted. Unfortunately, both conditions are hardly satisfied in clinical practice, especially when there is still no effective treatment to effectively halt the intrinsic deterioration of BF. The estimated time when BF deterioration reaches 50% of its maximum shows a high variability, ranging from 0.0203 ∼ 4.10 years, with an average of 0.215 years, implying that β-cell deterioration could reach its maximum rate within only a few months if without any intervention. The patients with the lowest estimate of *DP*_*50*_ (0.0203 years) were those whose β-cells had almost completely lost the restoration potential at diagnosis. Once the deterioration rate reaches its maximum, the major factor that influences BF deterioration is glucotoxicity and strictly controlling glucose would be critical to long-term management of T2DM.

The parameters were also compared across intervention groups. As shown in Table 1, sulfonylureas had the fastest rate of β-cell deterioration (*k*_*out_beta*_), suggesting a limited protective effect on β-cells. The conventional treatment (exercise and diet) reached 50% of the maximum deterioration in the shortest duration (DP_50_), indicating that conventional treatment did not have as much protective effect on β-cell as the therapeutic agents.

The ratio of *k*_*in_beta*_/*k*_*out_beta*_ indicates the theoretical potential of β-cell restoration when PFG is strictly controlled within the normal level (4.5 mM). As shown in **Fig. 4**, the theoretical potentials of BF restoration after interventions are correlated to the initial BF at diagnosis. This sigmoidal correlation indicates that a higher BF at diagnosis predicted a higher BF restoration potential. On average, β-cells can restore 80% of their normal function for early-stage diabetes. Even though such restorations are only temporary, they have long-term benefits in diabetes control. Fig. 4 shows that there was no apparent BF restoration when the initial BF was below 40% (potential BF < initial BF), no matter what type of therapeutics were adopted and how strictly the glucose was controlled. These observations strongly suggest the importance of early diagnosis and early intervention for T2DM.

**Fig. 4.**
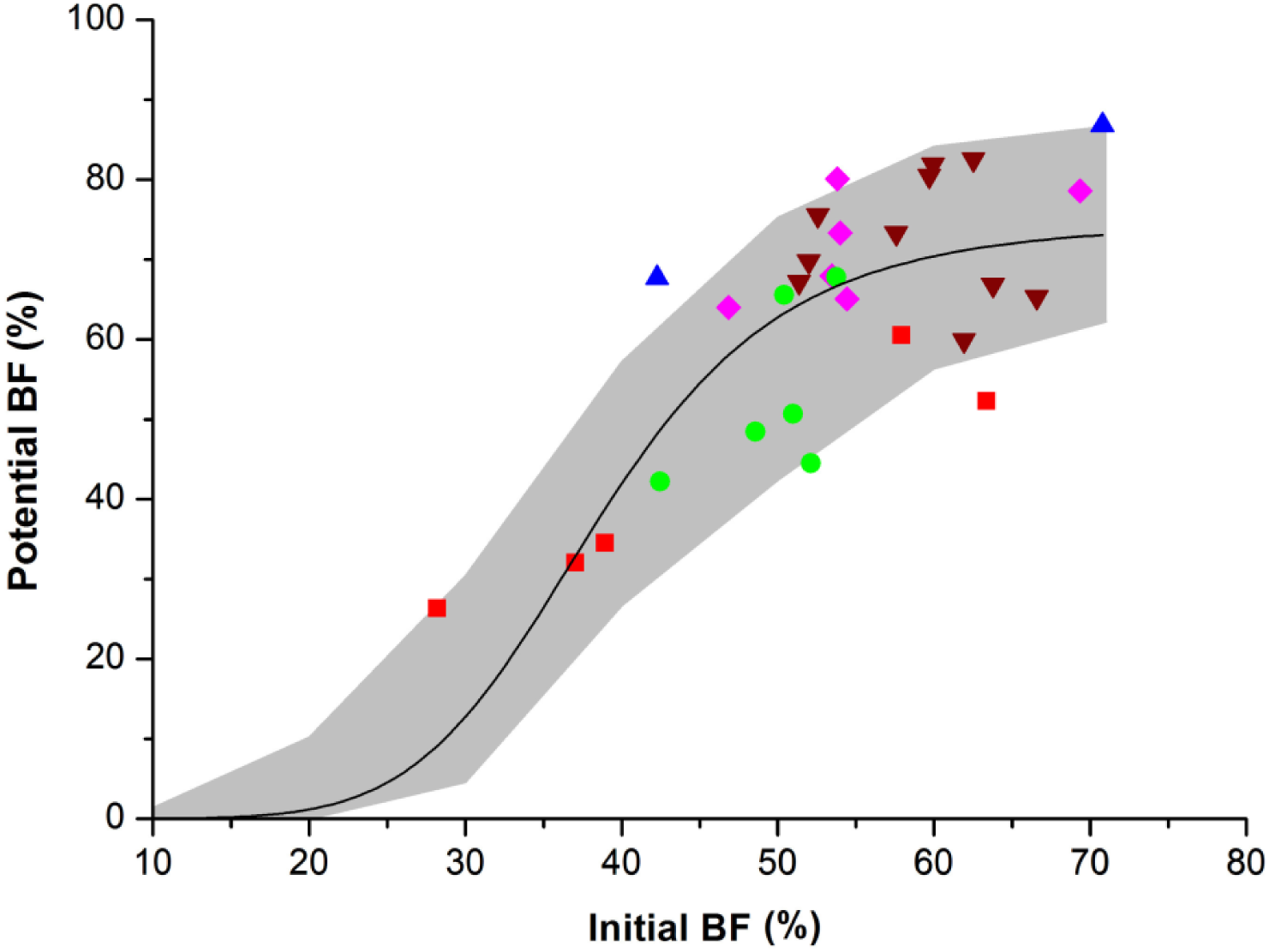
Relevance between first BF and potential BF (*k*_*in_beta*_/*k*_*out_beta*_). Legend: pink diamond = conventional group; green circle = insulin group; brown triangle = sulfonylureas group; red square = metformin group; blue triangle = thiazolidinediones group; solid lines = the simulated curve for all real observations by sigmoid function; gray shaded areas = between the 10th and 90th percentiles of the simulated data.

As shown in **Fig. 5**, the BF profiles were simulated under three plasma glucose levels (4.5, 8.3, and 11.1 mM). As expected, PFG around 4.5 mM yielded a considerable β-cell restoration, while a PFG higher than 11.1 mM completely eradicated BF restoration. The PFG threshold for BF restorations was predicted to be approximately 8.3 mM when the initial BF is 55% at diagnosis. The strong effect of glucose control on BF restoration supports our current therapeutic philosophy of T2DM, which is to initiate insulin therapy as early as possible to strictly control the glucose level, even at a very early stage of T2DM, for better BF protection and long-term glycemic management (Weng J et al., 2008).

**Fig. 5.**
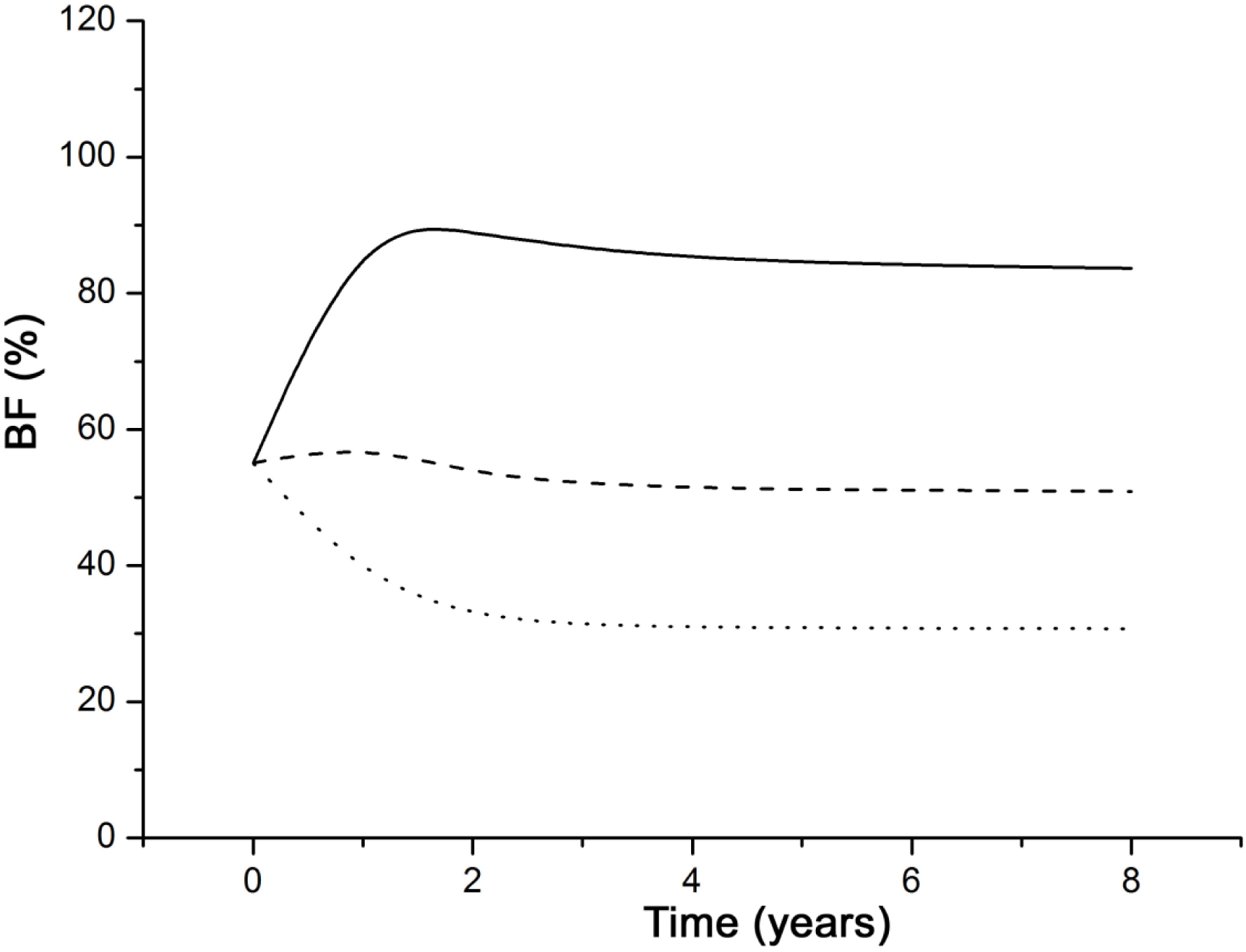
Influence of hyperglycemia on BF in three given conditions. The solid line indicates initial PFG=4.5 mM; the dashed line indicates initial PFG=8.3mM; the dotted line indicates initial PFG=11.1 mM.

As shown in **Fig. 6**, the long-term profiles of BF and PFG were simulated as a function of the β-cell intrinsic deterioration rate (*DP*_*50*_ = 0.1, 0.5, or 1.0 year). As noted, a several-month delay in BF deterioration at an early stage yielded a significant improvement in long-term glycemic control and BF preservation. The probability of BF below 50 % decreased from 66.9 % to 51.1 % when prolonging *DP*_*50*_ from 0.1 to 1.0 year, and the probability of PFG over 10 mM at 8 years of diagnosis was reduced from 31.7 % to 22.5 %. Although the β-cell deterioration rate reaches its maximum shortly after diagnosis, a moderate delay of β-cell deterioration at an early stage was predicted to have a long-term benefit.

**Fig. 6.**
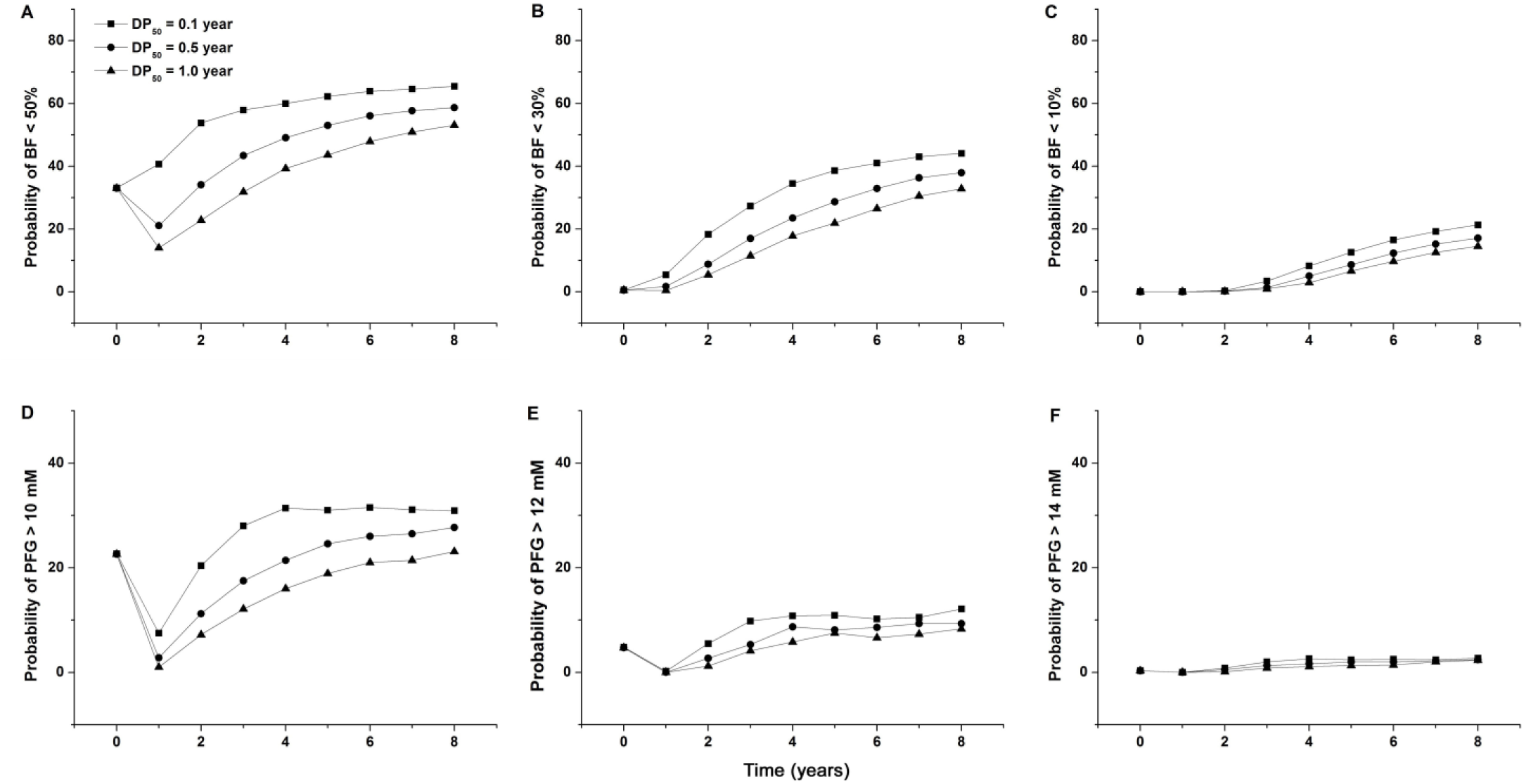
Influence of β-cell deterioration rate on BF and PFG over eight years. Probability was based on 1000 simulations.

**Fig. 7** summarizes the probability of BF and PFG higher or lower than designated levels when insulin resistance is improved (ISI = 0.15, 0.18, 0.20). The probability of PFG levels above 10 mM after eight years of diagnosis is 52.1 %, 39.4 %, and 31.0 % when the ISI is 0.15, 0.18, and 0.20, respectively. The probabilities of PFG over 12 mM vs over 14 mM also showed significant differences with a moderate change in insulin sensitivity. The probabilities of BF levels lower than 10 %, 30 %, or 50 % were also largely different. The substantial enhancement in long-term glycemic control and BF restoration with a moderate improvement of ISI (from 0.15 to 0.20) highlights the importance of re-sensitizing the insulin response in T2DM management.

**Fig. 7.**
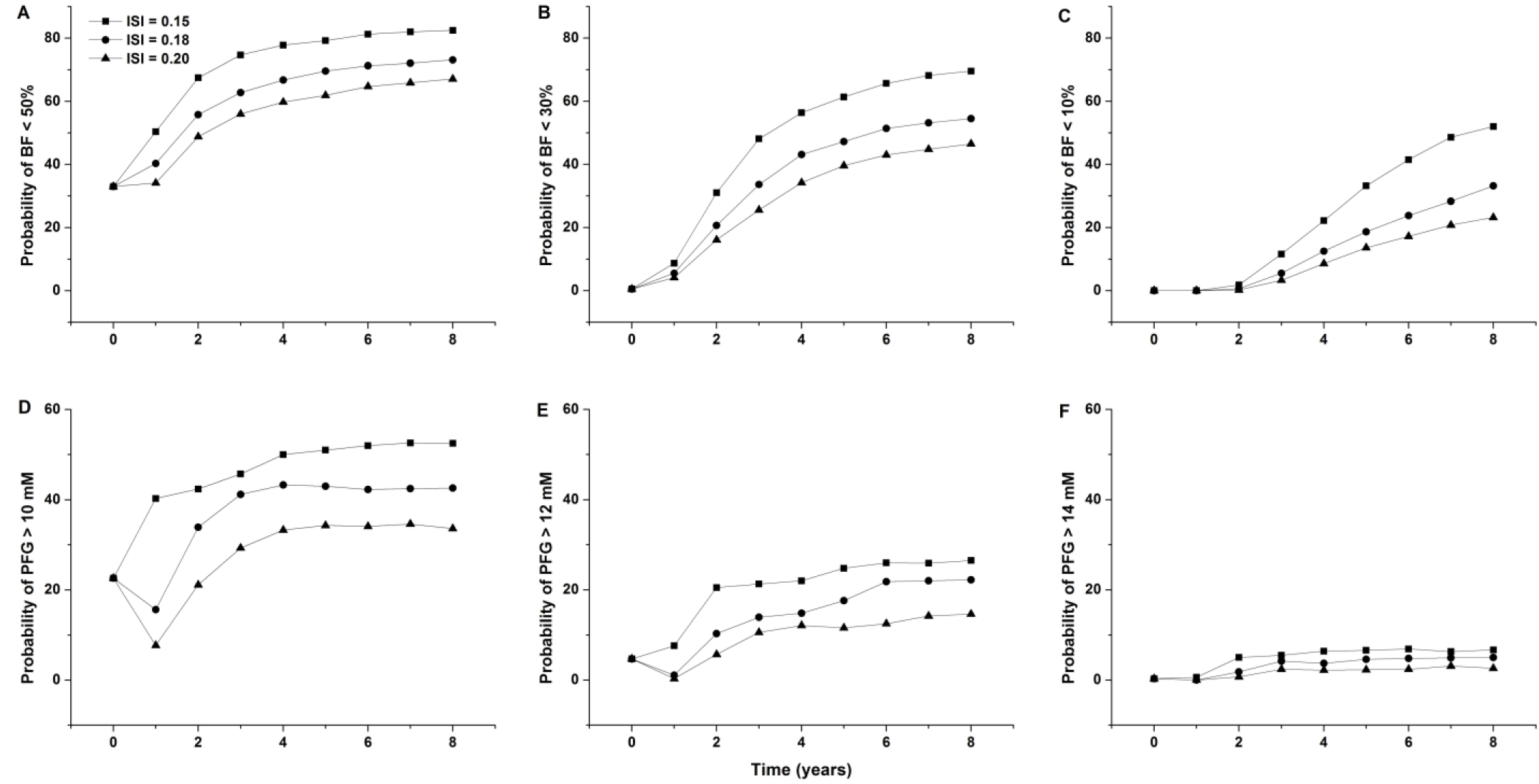
Influence of insulin resistance on BF and PFG over eight years. Probability was based on 1000 simulations.

## 4. Discussion

A mechanistic disease progression model for T2DM was developed in this study. The model used data extracted from multiple clinical trials of long-term T2DM treatments. It integrated a dynamic interaction between BF and PFG and a natural deterioration process in BF. The model adequately captured both the trend and variability of BF and PFG over a long period starting from diagnosis. Several factors were simulated and compared to investigate effective treatment strategies for the long-term management of T2DM.

The present disease progression model was built based on HOMA-1 model estimates of BF and ISI. The HOMA model was developed initially by Matthews and co-workers for diabetes diagnostic purposes (Matthews DR et al., 1985). HOMA-1 is a theoretical approximation that is adjusted to population norms (Jevy JC et al., 1998). The HOMA model uses PFG and FI to estimate the remaining BF and ISI values in healthy subjects and T2DM. The model has been validated using a euglycemic clamp method (Matthews DR et al., 1985). During long-term clinical trials, the HOMA model provides a simple way to evaluate diabetic status, mainly when frequent glucose tolerance tests or euglycemic clamp assessments are not readily feasible. In addition, the model considers glucotoxicity on BF and a natural deterioration process of BF for a long-term characterization of diabetic progression in newly diagnosed T2DM.

Several diabetes disease models were reported in previous literature (Finegood DT et al., 1995; Landersdorfer CB and Jusko WJ, 2008). Topp et al. developed a disease model describing both the stimulating and detrimental effects of hyperglycemia on BF, in which the intrinsic deterioration of BF was not considered, and the model would not predict the restoration of BF after the intervention (Topp B et al., 2000). Bagust et al. described a β-cell deterioration process using a biphasic equation; the model did not include glucose dynamics and would not predict the BF restoration either (Bagust A and Beale S, 2003). De Winter used an asymptotical equation to mimic the deterioration process of BF, which was further used to investigate several anti-diabetes agents. The model did not quantitatively consider glucotoxicity so that it would not predict BF restoration (De Winter W et al., 2006). In terms of glucotoxicity, a study has investigated the quantitative relationships between PFG and BF, and a hyperbolic curve was previously indicated between PFG and beta-cell volume (Ritzel RA et al., 2006). The equations for glucotoxicity, including linear, nonlinear, or exponential relationships, were assessed in our model, and a linear relationship was selected, which was further proven to capture the data quite well. A logistic model was originally tested to describe the deterioration kinetics of BF, which did not converge successfully. In the present model, a time-dependent nonlinear model was thus selected, which was shown to be consistent with the observations.

The maximum tolerated level for PFG that supports BF restoration in early diabetes was suggested to be around 8.3 mM, which is higher than the American Diabetes Association’s recommended boundary of 7.2 mM (Nathan DM et al., 2006). Our results support the notion that early treatment in the pre-diabetes period is critical even if plasma glucose remains in the normal range at this stage. Early intervention anticipates a high restoration potential in BF. Treatments to improve BF, rather than solely controlling hyperglycemia, should be more productive during this treatment period for long-term benefits. Although β-cell-intrinsic deterioration occurred quickly following diagnosis, a slight delay in this process could drastically improve long-term glycemic control and preserve BF.

Although our model offers a satisfactory characterization of diabetes progression in early T2DM, a couple of limitations need to be clarified. First, BF and ISI were derived from the HOMA-1 model, which made it improbable to separate the treatment effects from the underlying disease status. For example, the use of insulin-secreting drugs could boost insulin concentrations and thereby bias the calculation of BF related to the actual insulin-producing capacity of the β-cells. Therefore, the calculated values for BF and ISI were confounded by the intervention effects. Another limitation is related to the empirical approach in which we formulated glucotoxicity in equation 3. The quantitative glucotoxicity on BF is still unclear, and more quantitative analysis is warranted to clarify Equation 3.

## 5. Conclusion

The developed diabetes model integrated β-cell intrinsic deterioration, glucotoxicity, insulin resistance, as well as their dynamic interactions to depict diabetes progression over eight years. The model sufficiently predicted the long-term profiles of BF and PFG, highlighting the importance of early diagnosis and early interventions in T2DM for a long-term benefit to glucose control.

## Abbreviations

BF: β-cell function
FI: Fasting insulin
HOMA-1: Homeostasis model assessment 1
HOMA-2: Homeostasis model assessment 2
ISI: Insulin sensitivity index
PFG: Plasma fasting glucose
T2DM: type 2 diabetes mellitus

## Acknowledgements

This work was supported by National Institute of Health (GM119661).

## Notes

### Competing Interest Statement

The authors have declared no competing interest.

